# Molecular biological methods to assess different *Botrytis cinerea* strains on grapes

**DOI:** 10.1101/2024.01.12.575343

**Authors:** Louis Backmann, Katharina Schmidtmann, Pascal Wegmann-Herr, Andreas Jürgens, Maren Scharfenberger-Schmeer

**Affiliations:** Institute for Viticulture and Oenology, Dienstleistungszentrum Ländlicher Raum (DLR) Rheinpfalz, Breitenweg 71, D-67435 Neustadt, Germany; Hochschule Kaiserslautern, Weincampus Neustadt, Breitenweg 71, D-67435 Neustadt, Germany; Department of Biology, Chemical Plant Ecology, Technische Universität Darmstadt, Schnittspahnstrasse 4, D- 64287 Darmstadt, Germany

**Keywords:** *Botrytis cinerea*, strain differentiation, simple sequence repeat markers, qPCR, early detection

## Abstract

*Botrytis cinerea* is a well-known pathogen that can be challenging to control in crops, such as wine grapes. To adapt to the increasing problems of climate change and strain resistance it is important to find new methods to detect *Botrytis cinerea* and differentiate strains. These methods include strain differentiation and classification by simple sequence repeats (SSRs) and early detection of the fungus by qPCR. Various strains were analysed using SSR markers and either agarose gel electrophoresis or capillary sequencing via PCR. A sensitive qPCR method was refined to achieve an early detection method for the pathogen. The results demonstrate promising ways to distinguish between strains using both agarose gel electrophoresis and capillary sequencing as well as to detect infection before it becomes visible on grapes. This can be used to further understand and analyse different *Botrytis cinerea* strain characteristics such as laccase activity, regional or annual effects. The early detection method can be used to better prepare growers for an impending infection so that targeted efforts can be made.

## 1 Introduction

*Botrytis cinerea* spp. is a well-known pathogen that can cause crop failure. It infects over 200 different types of crops [1-2] including tomatoes, strawberries and grapes. Other related species such as *Botrytis allii, Botrytis byssoides, Botrytis squamosa, Botrytis fabae* and *Botrytis gladioli* are pathogens in onions, beans and flowers such as gladioli [3]. *B. cinerea* is a fungus that is found globally, particularly in cold and humid climates [4], as well as in temperate and subtropical climates [5]. It infects grapes (*Vitis vinifera*) and causes significant crop losses such as grey mould or bunch rot, resulting in economic losses of $ - 10-100 billion per year [6]. *Botrytis* enters grapes through damaged tissues during grape development and remains unnoticed until the grape matures further. This leads to rapid tissue decay in a short period of time, resulting in the harvesting of infected grapes [1]. Additionally, it promotes secondary infection with other pathogens such as *Penicillium expansum* [7]. Infection results in the production of laccase [8-9], a member of the blue copper oxidases [3]. Laccase oxidizes polyphenols into quinones, which then polymerize into brown compounds [10-11]. This process alters the colour of must and wine [12], leading to wine instability and colour degradation. Off-flavours such as geosmin or 1-octen-3-one [13-14] can affect wine quality at concentrations as low as 5% of *Botrytis* infected grapes [15]. These compounds result in earthy and mushroom-like flavours. The studies by La Guerche et al. [13-14] have also shown their impact on wine quality. *B. cinerea* growth not only affects the fermentation process in winemaking, but also has a negative impact on wine production [16]. Therefore, it is important to consider prevention measures rather than just treatment. Although there are numerous scientific studies and strategies to minimise the impact of *B. cinerea*, such as applying chemical and biological fungicides on grapes [17] or adding of bentonite or oenological tannins to must and wine [12], controlling the disease is challenging due to various attack pathways and survival strategies of *Botrytis* [6]. These studies have identified a diverse range of potential hosts and both sexual and asexual forms of survival [6, 18-20]. Furthermore, ongoing climate change resulting in warmer climates and extreme weather conditions has led to the evolution of more aggressive strains [21]. This has resulted in higher crop losses in wine, even in hot years [22], indicating the problematic future associated with *Botrytis*.

There are two main strategies to combat *B. cinerea*. The first strategy involves applying different treatments to prevent the growth of *Botrytis*, such as botricide, or to reduce the impact of the fungus on the must and wine by using oenological treatments such as active coal, tannins, or flash pasteurisation. To reduce harvest losses and improve the treatment of different *Botrytis* strains it is important to adapt treatments according to their environmental impact and classify them accordingly. It is crucial to reduce the use of fungicides and other treatments in the wine industry as outlined in the European Green Deal [23] due to the potential risks they pose to human health and environment [24]. Additionally, *B. cinerea* has demonstrated resistance to several fungicides [25-27]. Microbiological methods, such as SSR-PCR or qPCR, have been shown to be promising tools for gaining a better understanding of the pathogen. For example, Fournier et al. [28-29] developed an SSR-PCR assay to distinguish between different *B. cinerea* strains and to find differences between noble rot and grey mould [29].

The second strategy is to detect *Botrytis* infection at an early stage, before visible signs appear on the grapes. This approach enables more specific and efficient control of the fungus, before the rapid growth of *B. cinerea* causes significant damage. To achieve this, the fungus biomass can be quantified using a highly sensitive method such as qPCR. Quantification of *B. cinerea* by qPCR has already been explored [30]. However, the issue of cross-contamination and the potential impact of different *B. cinerea* strains on qPCR results has not yet been investigated.

This study investigates methods to differentiate between various strains of *B. cinerea*, based on the techniques used by Fournier et al. [28]. Additionally, we evaluate detection methods for quantifying *B. cinerea* in grapes using qPCR, considering the impact of different *B. cinerea* strains and cross-contamination with other pathogen strains present on grapes. The overall objective is to gain a better understanding of the diversity and distribution of *B. cinerea* strains.

## 2 Materials and Methods

### 2.1 Material

#### 2.1.1 Strain sampling and cultivation

During the 2022 harvest season, various *Botrytis cinerea* strains were obtained from different local regions and grape varieties including Rheinpfalz, Pfalz, Rheinhessen, Bonn, Elsass, Bordeaux, Baden, Freiburg im Breisgau. Additionally, strains from the DLR RLP Phytomedicine collection from previous harvest seasons (Pfalz) and one DSMZ Strain 877 were used. The strains were cultivated on 2 % Biomalz Agar (BA) plates at 25 °C and refreshed every 2-4 weeks. *Yarrowia Lipolytica* yeast was cultured on potato dextrose agar (PDA) plates and incubated at 30 °C for two days. *Cladosporium spp*., *Trichothecium roseum* and *Penicillium expansum* were grown on BA.

#### 2.1.2 Preparation of field samples

Grape samples of the Pinot Noir and Riesling varieties were obtained at different times during the harvest season. A total of 50 grapes were collected throughout the vineyard. An average sample of the vineyard was obtained by counting and packing 3 sets of 100 berries into separate bags, which were then stored in the freezer. The berries were crushed, using an Ultra-thorax (MICCRA, Buggingen, Germany) to homogenize the sample prior to use. For DNA extraction, 0,5 g of each sample was centrifuged at 13.000 g for 15 minutes and the resulting pellet was retained. Subsequently, 8 × 106 cells of *Yarrowia Lipolytica* were added to the sample, as an internal positive control for qPCR, and the mixture was centrifuged again for 15 minutes at 13.000 g. The resulting pellet was retained and subject to extraction following the RED Extract Plant PCR-Kit (Merck KGaA, Darmstadt, Germany) protocol.

### 2.2 Methods

#### 2.2.1 DNA Extraction

The strains and samples were extracted using the RED Extract Plant PCR kit from Merck KGaA, Darmstadt, Germany. Pre-tests have shown that this kit is an effective tool for quickly obtaining fungal DNA. Specifically, 100 µL of extraction solution was added to the prepared sample, mixed by vortexing, and incubated in a heat block at 95 °C for 10 min. Subsequently, 100 µL of dilution solution was added mixed by vortexing the resulting mixture was used for further analysis. After centrifugation at 13.000 g for 10 minutes, the soluble phase was retained and stored in the freezer for future use.

#### 2.2.2 qPCR

##### 2.2.2.1 Preparation of cultivated Botrytis samples – cross contamination

*Botrytis cinerea* strains were grown to promote sporulation and then removed from the plates. *Penicillium expansum, Trichothecium roseum* and *Cladosporium spp*. were also grown to promote sporulation and added separately to the *B. cinerea* probes. DNA was extracted using the method described above.

##### 2.2.2.2 Preparation of standard curves

During the experiment, various strains of *B. cinerea* were used to create the standard curve (Table 1). To create the qPCR standard curve for *B. cinerea*, a solution of 107 spores/ml was obtained by filtering *B. cinerea* mycel from agar plates after sporulation through a sieve and adding water. The spore count was determined using a light microscope and Neubauer counting chamber, and a solution of 107 cells/ml was created. Serial dilutions were used to obtain spore solutions ranging from 102 to 107 cells/ml. DNA was extracted using the RED Extract protocol. The *Y. Lipolytica* standard curve was obtained in a similar manner to the *B. cinerea* standard curve, with the addition of using an additional 108 cells/ml solution.

**Table 1.**
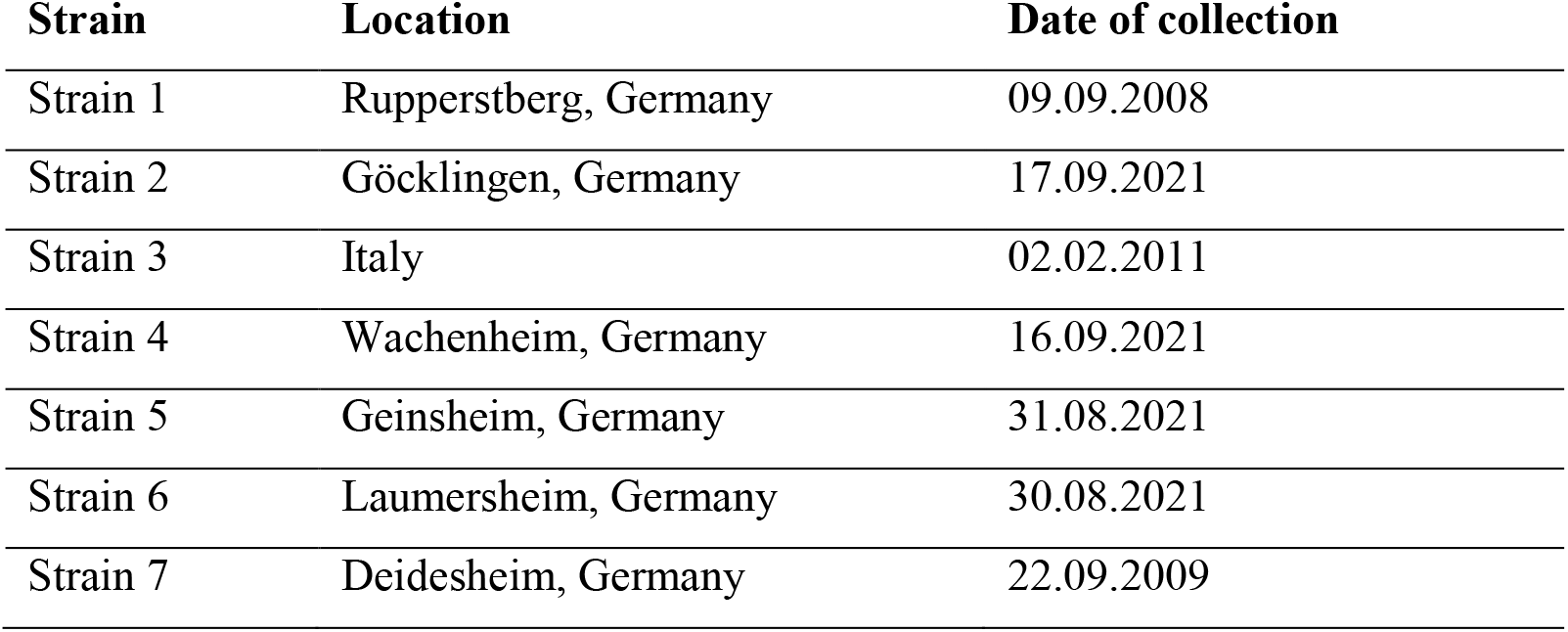
*Botrytis cinerea* strains used for qPCR and their location and date of collection.

| Strain | Location | Date of collection |
| --- | --- | --- |
| Strain 1 | Rupperstberg, Germany | 09.09.2008 |
| Strain 2 | Göcklingen, Germany | 17.09.2021 |
| Strain 3 | Italy | 02.02.2011 |
| Strain 4 | Wachenheim, Germany | 16.09.2021 |
| Strain 5 | Geinsheim, Germany | 31.08.2021 |
| Strain 6 | Laumersheim, Germany | 30.08.2021 |
| Strain 7 | Deidesheim, Germany | 22.09.2009 |

##### 2.2.2.3 qPCR run

The adapted version of the primer (Table 2) was used to measure all probes and standard curves in triplets, following the protocol described in Diguta et al. [30]. The qPCR protocol consisted of a hold stage at 95 °C for 3 minutes, followed by a 2-step PCR (95 °C 15 seconds, 65 °C 30 seconds – 40cycles), melting curve (90 °C 15 seconds, 50 °C 60 seconds, 95 °C 1 seconds). The resulting ct-values of the probes were analysed using the qPCR standard curves.

**Table 2.**
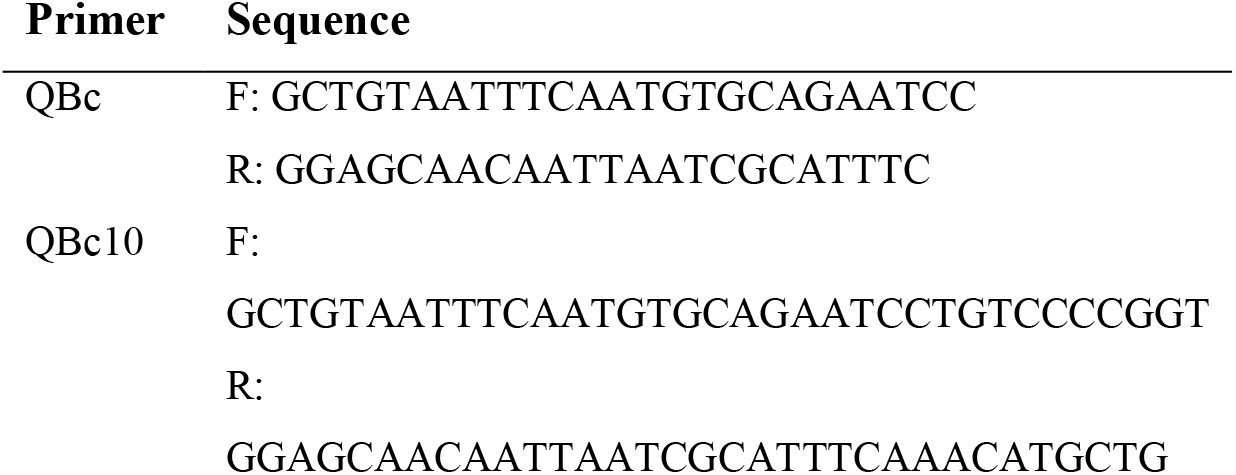
*Botrytis cinerea* Targeting Seq. Ribosomal Region 28S, 18S (Suarez et al., [31])

##### 2.2.2.4 qPCR as an early detection method - Limit of Detection

In the experiment, 300 berries per variant (Thompson Seedless) were used. A control with 0 spores/berry was compared to a variant with 10,000 spores/berry. The berries were “surface-sterilized” for 30 seconds in a 70% ethanol washing solution. Subsequently, a 10 µL droplet with 10,000 spores and a 10 µL droplet containing water (control) were applied to each berry. Every 24 hours, 3 × 10 berries were collected per variant, extracted and qPCR was performed as previously described. The experiment was continued until first sporulation was visible.

#### 2.2.3 SSR-PCR

##### 2.2.3.1 PCR run

A total of eight different simple sequence repeat markers, established by Fournier et. al [28] were used to perform PCR. The primers were tested for their size range at different temperatures and in different pairs to obtain optimal sets for a multiplex set of primers. The resulting multiplex sets were 1,2,4 and 3,5,6 at 50 °C and 7,10 at 60 °C (Table 3). PCR was performed following a standard procedure: The reaction tubes were prepared by adding 10 µL of RED Extract (Sigma-Aldrich), 4 µL MilliQ water, 1µL Forward/Reverse Primer each, and 4 µL sample DNA. The PCR program consisted of 36 cycles of denaturation at 94 °C for 30 seconds, annealing at primer dependent temperature (50 °C, 60 °C) for 30 seconds, and elongation at 72 °C for 30 seconds for a total of 36 cycles, preceded by initial denaturation at 94 °C for 3 minutes and followed by a final elongation at 72 °C for 10 minutes. The PCR was performed using an Eppendorf MasterCycler personal 5332 (Eppendorf SE, Hamburg, Germany). The PCR samples were stored at -20 °C in the freezer or at 4 °C in the fridge for later use.

**Table 3.**
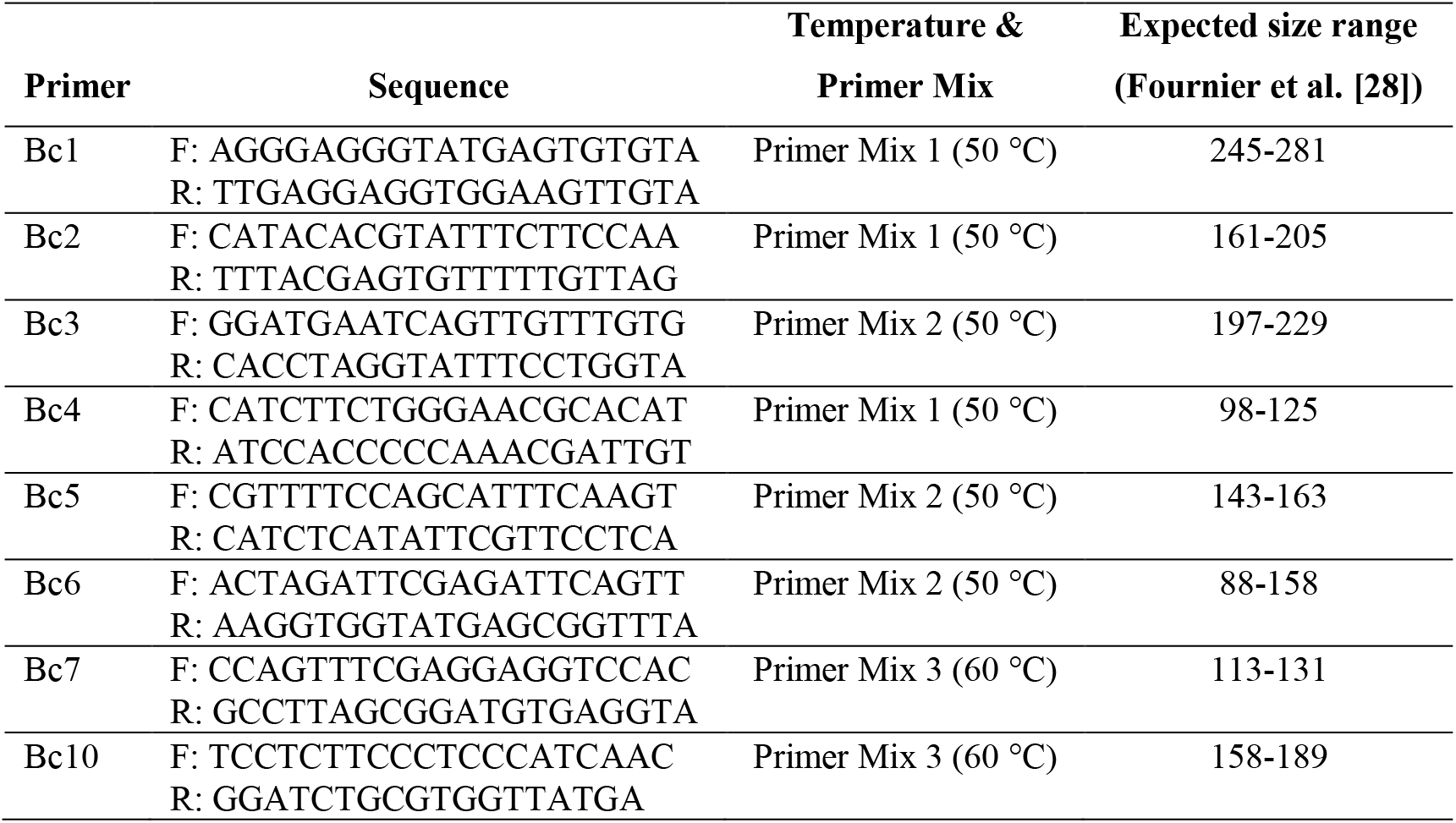
Primers sequence and multiplex composure and the expected size range of the resulting products (basepairs)

##### 2.2.3.2 Agarose Gel electrophoresis

To perform agarose gel electrophoresis, 3.1 g agarose (AppliChem GmbH, Darmstadt, Germany) were added to 100 ml of 10X TAE buffer (AppliChem GmbH, Darmstadt, Germany) and microwaved at 600 W for 3 to 4 minutes until the mixture became smooth. After cooling the mixture to -50 °C, 10 µL GelRed (GeneON, Ludwigshafen, Germany) was added. The gel was prepared by adding 5 µL of the PCR probes and 2 µL of the ladder (Thermo Fisher Scientific, Waltham, MA, USA) and running the reaction at 90 V for 2 hours and 45 minutes. The resulting gel was captured using a UV camera lens.

##### 2.2.3.3 Evaluation of agarose Gel

To assess the resulting gel bands, we measured the bands on the captured images using the ImageJ software tool [32] and compared them with the ladder. We compared resulting PCR product sizes to obtain a set of band sizes for each tested strain.

##### 2.2.3.4 Capillary sequencer

A total of eight simple sequence repeat markers, established by Fournier et. al [28] were used to perform PCR. The primers were tested at different temperatures and in different pairs to obtain optimal sets for a multiplex set of primers. The obtained multiplex sets were Bc1, Bc5, Bc10; Bc2, Bc3, Bc6 and Bc4,

Bc7, Bc9 at 60 °C. The procedure followed the protocol by Huber et al. [33]. The KAPA2G Fast Multiplex PCR Kit (KAPABIOSYSTENS, USA) was used to conduct the multiplex PCR, which included up to 10 primer pairs with fluorescent labels (forward primer coupled with HEX, ROX, TAMRA or FAM, Table 4). The PCR program consisted of 30 cycles of denaturation, annealing and elongation, with initial denaturation at 95 °C for 3 minutes, denaturation at 95 °C for 15 seconds, primer annealing at 60 °C for 30 seconds, elongation at 72 °C for 30 or 50 seconds, and final elongation at 72 °C for 3 minutes. The length analyses of the fragments were performed using a 3130xl Genetic Analyzer (Applied Biosystems, Germany) and the corresponding GeneMapper 4.0. software. The PCR samples were stored at 4 °C in the fridge or in the freezer (−20 °C) for later use.

**Table 4.**
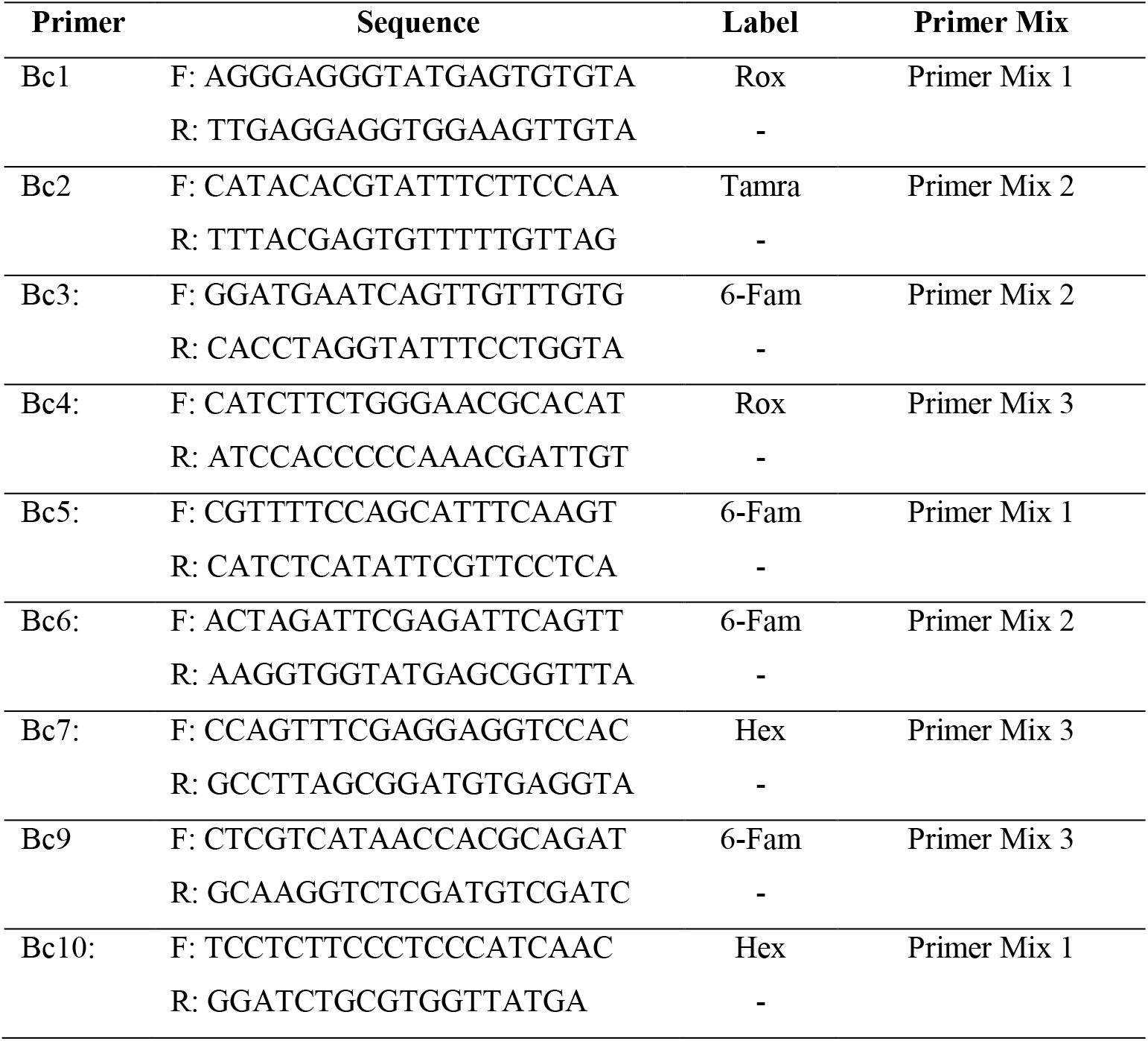
Table of the primers by Fournier et al. [28]:

## 3 Results

### 3.1 qPCR cross contamination

During an infection with *Botrytis cinerea*, cross-contaminations with different fungi can occur, which may affect the quantification of *Botrytis* biomass. To ensure primer specificity, the primers used in qPCR were tested against fungi that are commonly found on grapes including *Penicillium expansum, Trichothecium roseum* and *Cladosporium spp*.. *B. cinerea* and *P. expansum* were extracted separately and together and then tested for amplification using PCR and qPCR. The pure *Penicillium* probe did not show any bands in the PCR test, but qPCR showed a slight amplification of *Penicillium* using the original primer set. Therefore, the primer set was adapted by making the forward and reverse primers 10 bp longer on the 3’-end, according to the known genome sequence in Ensembl. Further testing revealed no amplification *Penicillium* or other tested fungi. Additionally, the primer exhibited no cross-reaction with the tested fungi (Figure 1). Both primer sets were free of primer dimers or by-products in the melting curve.

**Figure 1.**
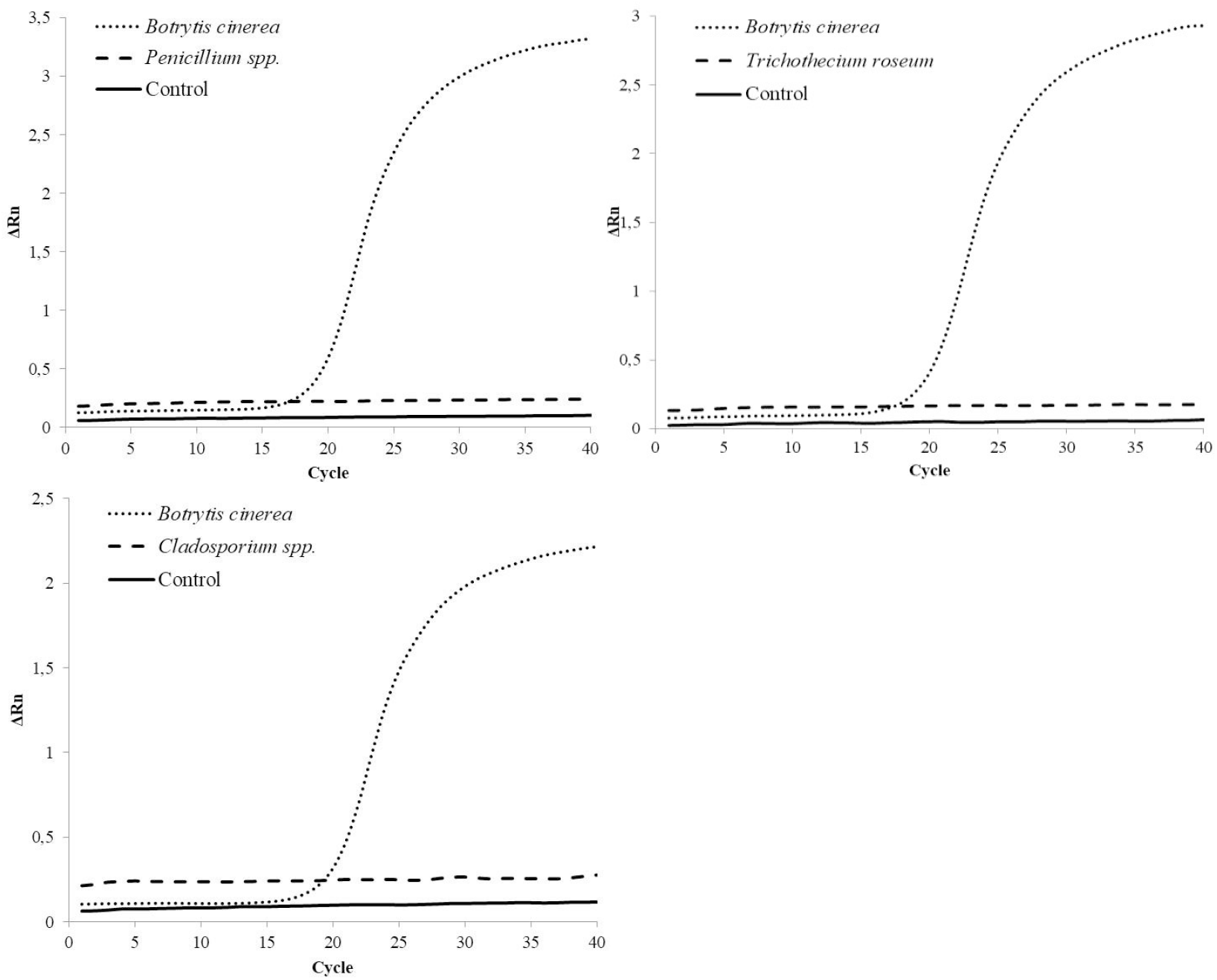
Cross contamination results of the QBc10 Primer. In every run the primer was tested against a *Botrytis cinerea* strain, either *Penicillium expansum, Trichothecium roseum* or *Cladosporium spp*. and a negative control.

### 3.2 Effect of different Botrytis strains on the standard calibration curve

To investigate the potential impact of different *B. cinerea* strains on qPCR efficiency, we prepared and counted spore suspensions under a light microscope to obtain both high and low concentrations of spores per millilitre. The strains used were: strain 1 (Ruppertsberg, Germany, 2008), strain 2 (Göcklingen, Germany, 2021), strain 3 (Italy, 2011), strain 4 (Wachenheim, Germany, 2021), strain 5 (Geinsheim, Germany, 2021), strain 6 (Laumersheim, Germany, 2021). The spore suspensions were extracted and quantified against a standard curve using the same *B. cinerea* strains for every measurement, specifically strain 7 from Deidesheim, Germany, 2009. The primer efficiency ranged from 94 % to 96 %. The results were then compared with the counted cell number as shown in Figure 2. Most of the *Botrytis* strains used for the standard calibration curve were within a reasonable variance range compared to the manual counting, considering the limitations of obtaining exact results using the counting chamber. However, strain 3 showed a greater variance of up to 23.5%.

**Figure 2.**
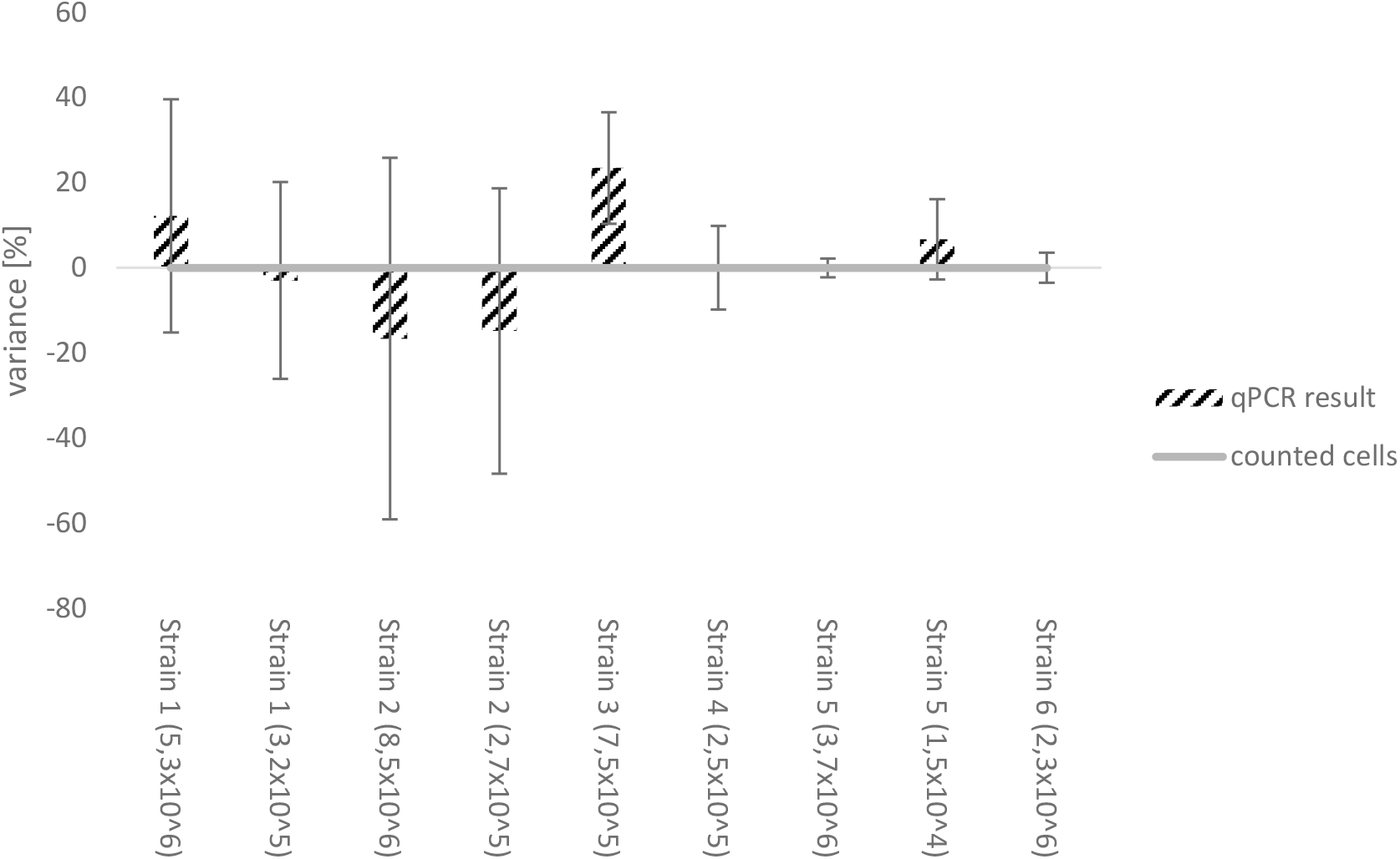
Comparison of the counted cell number to the different qPCR results using the same *Botrytis cinerea* strain as standard calibration curve (strain 7, Deidesheim, Germany, 2021). Strains used: strain 1 (Ruppertsberg, Germany, 2008), strain 2 (Göcklingen, Germany, 2021), strain 3 (Italy, 2011), strain 4 (Wachenheim, Germany, 2021), strain 5 (Geinsheim, Germany, 2021), strain 6 (Laumersheim, Germany, 2021) The results were normalized towards the counted cell number (Neubauer counting chamber). In some cases, two different cell concentrations representing a high and low cell number were analysed.

### 3.3 qPCR as an early detection method - Limit of Detection

In the first experiment infection levels were not reliable observed when using 100 and 1000 spores per berry. However, when using 10,000 spores per berry, infection levels above the detection limit were observed [34]. Therefore, 10,000 spores/berry were chosen for the next experiment. In the second experiment early signs of infection were observed on day 3 and 4, although no sporulation was visible on the berry. The detected biomass increased during the infection period and sporulation became visible after 7 days (see Figure 3). Paired t-tests were performed for every individual between the spore and control variant (*** = p<0,0001).

**Figure 3.**
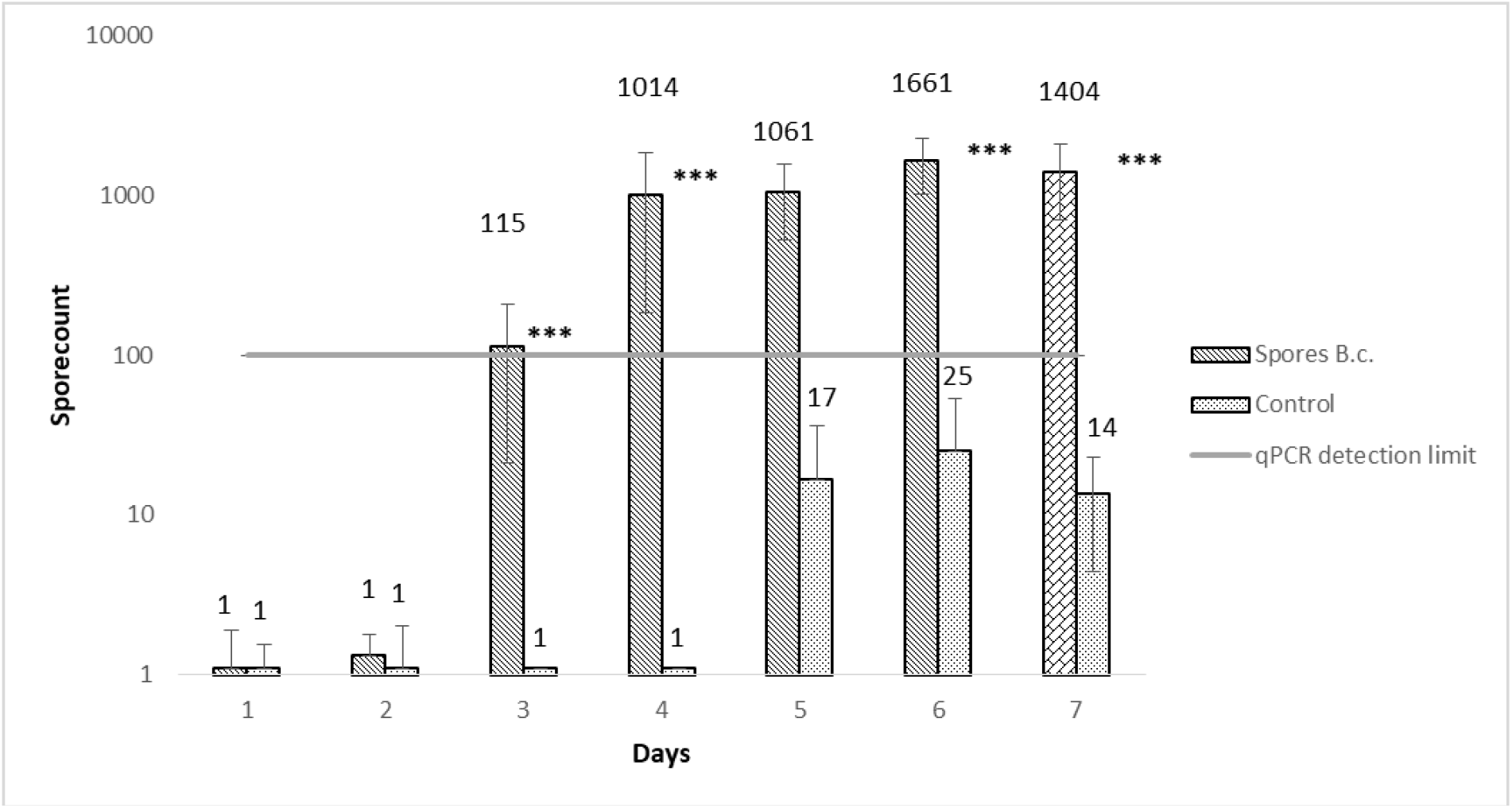
Early detection of *Botrytis cinerea* on table grapes. Berries were inoculated with 10.000 spores/berry and compared to a control. Every 24h 3x 10 berries were collected per variant, extracted and qPCR was performed. The experiment continued until sporulation was visible on the grapes (paired t-test; *** = p<0,0001)

### 3.4 PCR Method validation-Size ranges of Primer sets and composition

The amplified bands of the different simple sequence repeat markers were of varying sizes. A total of 70 strains underwent testing through capillary sequencer and agarose gel analysis and the results were compared (Table 5). Both methods were able to distinguish a large proportion of the tested strains. Sample images of the results from the capillary sequencer (Figure 4) and agarose gel analysis are presented in Figure 4 and Figure 5.

**Table 5.** Comparison of different methods to distinguish *Botrytis cinerea* strains using simple sequence repeat markers. In total, 70 strains were analysed by PCR followed by either an agarose gel electrophoresis or capillary sequencer. For every primer pair and method, the resulting basepair ranges of the strains are shown.

| Primer | capillary sequencer | agarose gel | Fournier et al. [28] |
| --- | --- | --- | --- |
| Bc1 | 219-265 | 195-256 | 245-281 |
| Bc2 | 144-183 | 132-180 | 161-205 |
| Bc3 | 213-221 | 189-240 | 197-229 |
| Bc4 | 118-127 | 107-132 | 98-125 |
| Bc5 | 117-162 | 135-169 | 143-163 |
| Bc6 | 109-125 | 105-130 | 88-158 |
| Bc7 | 111-119 | 105-120 | 113-131 |
| Bc9 | 146-147 | / | 150-194 |
| Bc10 | 178-189 | 165-200 | 158-189 |

**Figure 4.**
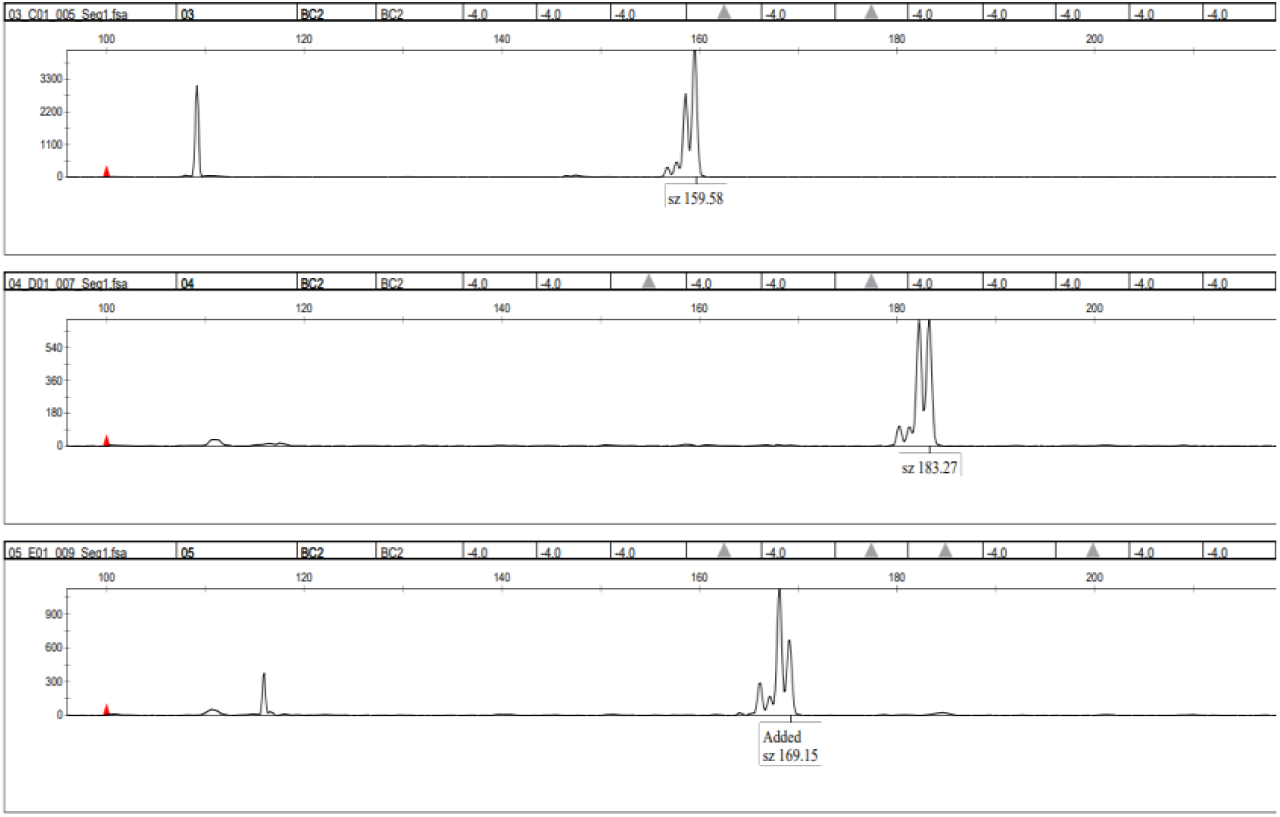
Sample picture of the capillary sequencer result. Labelled peaks in the diagram represent primer pair products in their respective basepair size. The tested strains were manually analysed after the experiment to ensure correct measurements of the software.

**Figure 5.**
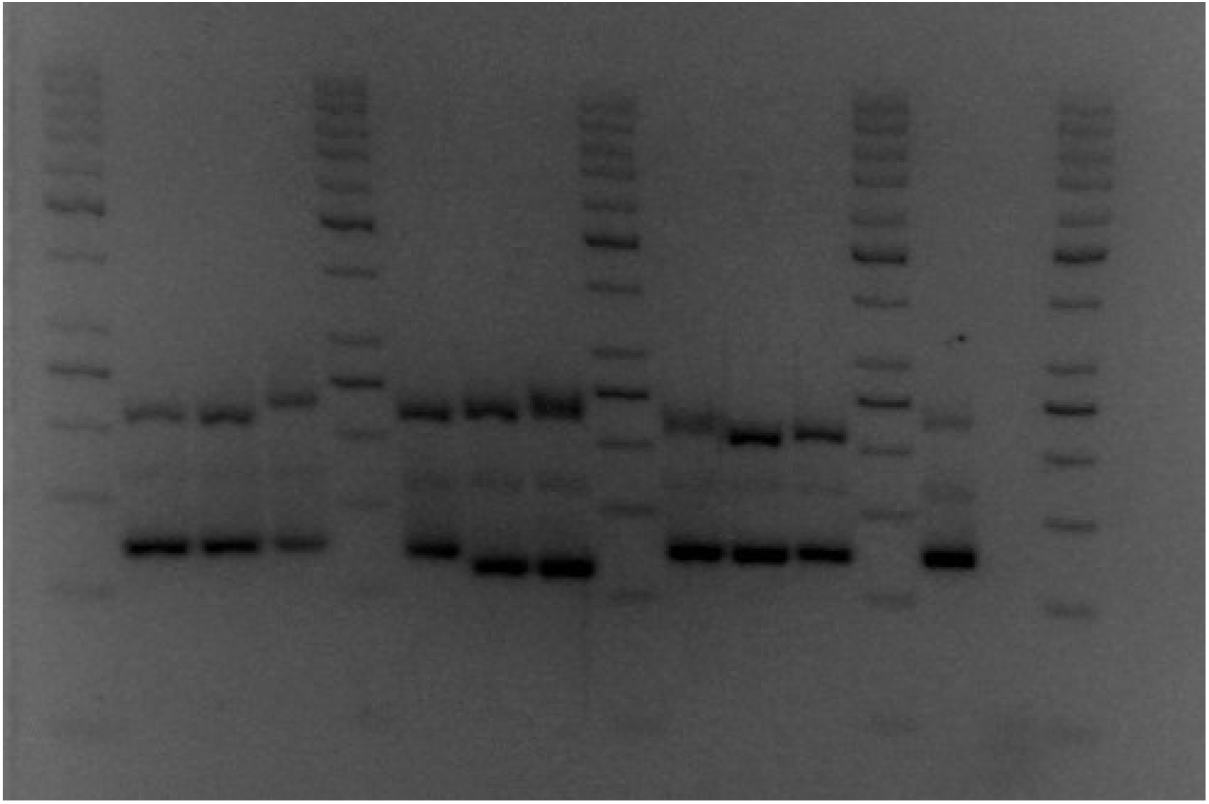
Sample picture of the Agarose gel electrophoresis. 10 *Botrytis cinerea* strains, one primer mix containing the primer pairs Bc1, Bc2 and Bc4. A normal run consisted of 10 strains and 1 negative control tested against 1 primer mix (either 1-2-4, 3-5-6 or 7-10). Runtime 2 h 45 m at 90 V.

## 4 Discussion

### 4.1 Cross contamination

The study of fungi that may occur as secondary infections on grapes after a *Botrytis cinerea* infection is crucial for obtaining accurate biomass quantification results. This is particularly important when analysing different strains with varying growth characteristics. The Bc10nt primer used for qPCR was validated for cross-contamination with the three fungi *Penicillium expansum, Trichothecium roseum* and *Cladosporium* spp. After modifying the Bc10 primer set by adding an additional 10 bp following the genome sequence, no cross-contamination was detected. This demonstrates that the primer can successfully quantify *B. cinerea* biomass in field samples. However, it should be noted that grapevines can also be affected by other pathogens, such as *Aspergillus niger*. However, analysing these organisms often requires a higher biosafety laboratory standard. Due to extended primer pair, the likelihood of cross-contamination should be low.

### 4.2 Effect of different Botrytis strains on the standard calibration curve

The qPCR analysis of various *B. cinerea* strains revealed only slight discrepancies between the number of spores counted and the number of spores quantified. For most strains, quantified biomass/spores were comparable to the manual cell count. The same strains were used as the qPCR standard curve when quantifying the biomass of the counted strains. The primer pair targets a highly conserved region in the genome. However, there are large differences between some of the strains used. This may be due to limitations in cell counting or differences in the amplified region targeted by the primer. Therefore, comparing a cultivated *Botrytis* strain in the standard curve with unknown field samples could lead to inaccurate results. It is not advisable to cultivate the potential *B. cinerea* strain of the field sample initially, as the qPCR results must be available quickly and should serve as an early detection method for winegrowers. However, qPCR can be used to compare different *B. cinerea* strains in the standard curve for research purposes or to analyse potential treatments where the season is not a factor. For early detection, any level of *B. cinerea* detection should serve as a warning sign to winegrowers. Exact quantitative results are not that important.

### 4.3 qPCR as an early detection method against Botrytis cinerea

When using qPCR, it is important to quantify the exact amount of biomass. Additionally, early detection of *B. cinerea* growth is crucial for monitoring pathogen growth in the vineyard. Measures can be taken to control the rot as soon as it is detected on the grapes. Currently, winegrowers rely on visual detection which can result in high crop losses if *Botrytis* is established. Winegrowers often attempt to predict potential infections based on prior knowledge and previous occurrences. However, this method is subjective and unreliable. The use of expensive botricides is often random. The implementation of qPCR to detect *B. cinerea* before the onset of rot would enable better preparation and more effective monitoring of the grapes, while reducing treatment costs. The qPCR analysis revealed that *B. cinerea* can be detected up to 3-4 days before visible infection on the grapes. Subsequently, there was an increase on day 6, with no further increase on day 7, when sporulation became visible. It is important to note that this experiment was conducted in a laboratory setting. Detection of infections in vineyards may be possible at an earlier stage if abiotic factors slow down the growth of *B. cinerea*. However, the infection process may be altered in table grapes due to their thicker skins compared to other grape varieties. Further research is needed to test the susceptibility of different classic grape varieties and fungus resistant grape varieties to *B. cinerea* infection.

### 4.4 PCR Method validation-Size ranges of Primer sets and composition

The simple sequence repeat markers amplified bands of the expected size (Table 5) with some variations from Fournier et al. [28]. Out of the 70 strains tested, both methods distinguished a significant proportion of the strains, with the capillary sequencing method distinguishing the most strains [34]. A typical agarose gel image can only differentiate the resulting bands of different strains up to 10-15 base pairs. Accurately distinguishing strains can be challenging due to issues with gel image resolution and band separation. To refine band separation, the gel run length can be adjusted, but if the run time is too long, the bands may become washed out. For unknown strains smaller bands may run out of the gel. Additional primers can increase selectivity. Capillary sequencers can determine PCR products with up to 1 bp accuracy. The capillary sequencer can distinguish more strains due to its higher resolution, compared to agarose gel analysis. This is because the primers used can be in overlapping base pair regions thanks to the use of fluorescent markers. As a result, more primer set combinations are possible with the capillary sequencer. However, not every laboratory has the capability to analyse strains with this device. Using a capillary sequencer is more expensive and requires more training than pouring and analysing an agarose gel. The choice of method should be based on the importance of strain differentiation, the laboratory’s budget and the operators’ experience. For instance, a possible solution is to use an agarose gel for routine work or pre-screening of strains, and then analyse the significant or indistinguishable strains with a capillary sequencer. The base pair range varied among the different primer pairs. Some primers exhibited low base pair variation, indicating a highly conserved target region. This demonstrates that not all primer pairs are equally important for strain differentiation. For instance, the majority of strains can be distinguished without the use primers Bc6 and Bc9, whereas primer pair Bc1 and Bc5 was able to differentiate most strains (Figure 6).

**Figure 6.**
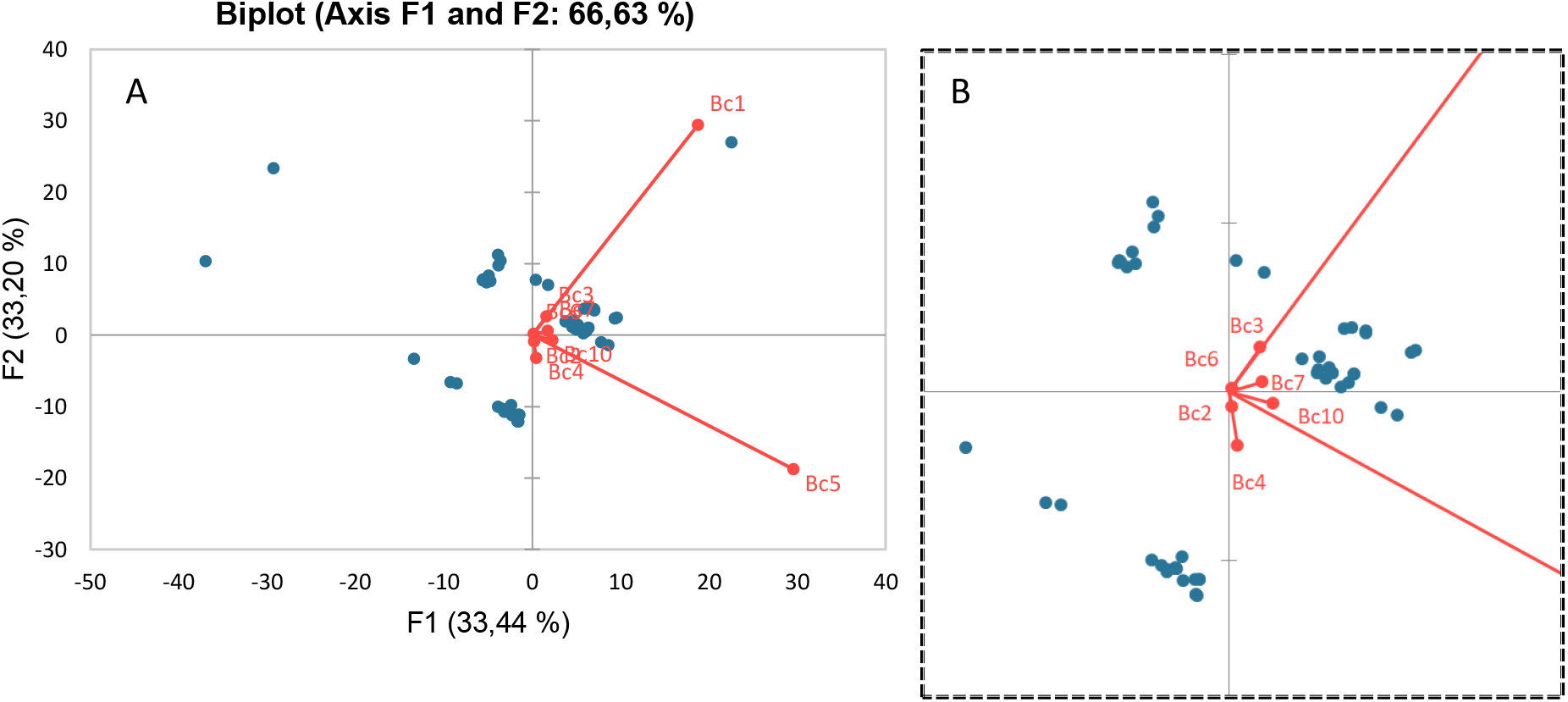
Principal component analysis (PCA) of the tested strains in SSR-PCR. The influence of the used primer pairs on the differentiation of the strains can be seen (A). Zoomed in perspective of the PCA (B).

## Conclusions

The use of Simple Sequence Repeat Markers demonstrates that there are various methodological approaches to detecting and analysing different strains of *Botrytis cinerea*. All of these approaches are valid, depending on available equipment and trained personnel. Further research could utilise different strains to investigate their effects on grapes, must and wine. It is important to consider the impact of climate change on the aggressiveness of different *B. cinerea* strains. Therefore, further research is necessary to determine whether traditional approaches to *B. cinerea* are still applicable. Strain differentiation can aid in identifying differences between *B. cinerea* strains based on location, region, grape variety, or time, which can enhance our understanding of *B. cinerea* on grapevines. The locations of the genes targeted by the primers could provide insights into the attack patterns and survivability of different strains. This information could be used to develop strain specific treatments, focusing on more aggressive strains.

The qPCR method was validated for quantifying *B. cinerea* on grapes by eliminating secondary infection quantification. This opens up more research potential for *B. cinerea* field studies. qPCR is a highly sensitive method that can detect a *B. cinerea* infection at an early stage before it is visually detectable. This could prove useful for winemakers to obtain accurate and early results of an infection in the vineyard. For instance, this could be used to improve the assessment of fungicide application and fungal monitoring, thus reducing potential treatment costs. An example of its potential application is the monitoring of vineyards for ice wine production, where grapes are even more susceptible to infection, and the loss of the entire harvest is possible. Quantifying potential undetected *B. cinerea* infection requires specialized laboratory equipment, making time and limit of detection the most important factors. Further research should be conducted to gain more insight into the detection method for *B. cinerea* infection, such as using different aggressive strains to determine the detection limit.

## Author Contributions

L.B. carried out the experiments. K.Z. assisted L.B. with the qPCR experiments regarding cross contamination and different strains for the standard calibration curve. L.B. wrote the manuscript with support from M.S-S. and A.J.. P.H. conceived the original idea. M.S-S. did the project administration and the funding acquisition. All authors have read and agreed to the published version of the manuscript.

## Acknowledgments

We acknowledge Florian Schwander and Margrit Daum for help with the analysis on the capillary sequencer.

## References

1. Williamson, B.: Tudzynski, B.: Tudzynski, P: Van Kan, J.A.L.: Botrytis cinerea: the cause of grey mould disease Molecular Plant Pathology 2007, 8(5), 561–580.

2. Andrew, M.: Barua, R.: Short, S.M.: Kohn, L.: Evidence for a common toolbox based on necrotrophy in a fungal lineage spanning necrotrophs, biotrophs, endophytes, host generalists and specialists. PLoS One 2012 7: 29943.

3. Claus, H.: Laccases of Botrytis cinerea. Biology of Microorganisms on Grapes, in Must and in Wine. H. König, G. Unden and J. Fröhlich. Cham, Springer International Publishing: 2017 339–356.

4. Steel, C.C.: Blackman, J.W.: Schmidtke, L.M.: Grapevine bunch rots: impacts on wine composition, quality, and potential procedures for the removal of wine faults.” J Agric Food Chem 2013 61(22): 5189–5206.

5. Bamford, D.H.: Zuckerman M.: Encyclopedia of Virology. Academic Press 2021 (4).

6. Poveda, J.: Barquero, M.: Gonzáles-Andrés, F.: Insight into the Microbiological Control Strategies against Botrytis cinerea Using Systemic Plant Resistance Activation.” Agronomy 2020 10(11).

7. Ribéreau-Gayon, P.: Glories, Y.: Maujean, A.: Dubourdieu, D.: Handbook of Enology – the Chemistry of Wine Stabilization and Treatments. Jon Wiley & Sons Ltd. 2006 (2).

8. Armijo, G.: Schlechter, R.: Agurto, M.: Munos, D.: Nunez, C.: Arce-Johnson, P.: Grapevine Pathogenic Microorganisms: Understanding Infection Strategies and Host Response Scenarios. Frontiers in Plant Science 2016 382(7).

9. Gilad, N.L.: Bar-Nun, N.: Mayer, A.M.: The possible function of the glucan sheath of Botrytis cinerea: effects on the distribution of enzyme activities. FEMS Microbiology Letters 2001 199 (1) 109–113.

10. Claus, H.: How to deal with Uninvited Guets in Wine: Copper and Copper-containing Oxidases. Fermentation 2020 6, 38.

11. Thurston, C.F.: The structure and function of fungal laccases. Microbiology 1994 140, 19–26.

12. Vignault, A.: Pascual, O.: Jourdes, M.: Moine, V.: Fermaud, M.: Roudet, J.: Canals, J.M.: Teissedre, P-L.: Zamora, F.: Impact of enological tannins on laccase activity.” Oeno One 2019 53(1).

13. La Guerche, S.: Chamont, S.: Blancard, D.: Dubourdieu. D.: Darriet, P.: Origin of (-)-geosmin on grapes: on the complementary action of two fungi, botrytis cinerea and penicillium expansum.” Antonie Van Leeuwenhoek 2005 88(2): 131–139.

14. La Guerche. S.: De Senneville, L.: Blancard, D.: Impact of the Botrytis cinerea strain and metabolism on (-)-geosmin production by Penicillium expansum in grape juice. Antonie van Leeuwenhoek 2007 92, 331–341.

15. Ky, I.: Lorrain, B.: Jourdes, M.: Pasquier, G.: Fermaud, M.: Gény, L.: Rey, P.: Doneche, B.: Teissedre, P.-L.: Assessment of grey mould (Botrytis cinerea) impact on phenolic and sensory quality of Bordeaux grapes, musts and wines for two consecutive vintages. Australian Society of Grape and Wine Research 2012 18, 215–226.

16. Marchal, R.: Salmon, T.: Gonzalez, R.: Kemp, B.: Vrigneau, C.: Willams, P.: Doco, T.: Impact of Botrytis cinerea Contamination on the Characteristics and Foamability of Yeast Macromolecules Released during the Alcoholic Fermentation of a Model Grape Juice.” Molecules 2020 25(3).

17. Lindemann, M.: Blaβ, W.: Botrytizide, in Böckler F.;Dill, RÖMPP [Online], Stuttgart, Georg Thieme Verlag, [Dezember 2023] (https://roempp.thieme.de/lexicon/RD-02-02370) 2007 RD-02-02370.

18. Caseys, C.: Shi, G.: Soltis, N.E.: Gwinner, R.: Corwin, J.A.: Atwell, S.: Kliebenstein, D.J.: Quantitative interactions drive Botrytis cinerea disease outcome across the plant kingdom. BioRxiv 2018 507491.

19. Mercier, J.: Roussel, D.: Charles, M.-T.: Arul, J.: Systemic and Local Responses Associated with UV- and Pathogen-Induced Resistance to Botrytis cinerea in Stored Carrot. Phytopathology 2000 (90) 981–986.

20. Mercier, A.: Carpentier, F.: Duplaix, C: Auger, A.: Pradier, J.M.: Viaud, M.: Gladieux, P.: Walker, A.S.: The polyphagous plant pathogenic fungus Botrytis cinerea encompasses host-specialized and generalist populations. Environmental Microbiology 2019 21(12) 4808–4821.

21. Shuping, D. S. S.: Eloff, J.N.: The Use of Plants to Protect Plants and Food against Fungal Pathogens: A Review. Afr J Tradit Complement Altern Med 2017 14(4): 120–127.

22. DLR Rheinpfalz 2020. Rebschutz 2020. Available online: https://www.dlr.rlp.de/Internet/global/themen.nsf/(Web_P_WB_Pfs_Kat_UKat_XP)/4AF73C1C0B641F94C12584FC003270A6/$FILE/Rebschutz%202020_03_Stand230320_AuflageY203.pdf:InstitutfürPhytomedizin. [Accessed 02.06. 2020].

23. European Commission. Available online: green deal https://commission.europa.eu/strategy-and-policy/priorities-2019-2024/european-green-deal_de (accessed 06.11.2023).

24. Hua, L.: Yong, C.: Zhanquan, Z.: Boqiang, L.: Guozheng, Q.: Shiping, T.: Pathogenic mechanisms and control strategies of Botrytis cinerea causing post-harvest decay in fruits and vegetables. Food Qual. Saf. 2018 2, 111–119.

25. Elad, Y.: Yunis, H.: Katan, T.: Multiple fungicide resistance to benzimidazoles, dicarboximides and diethofencarb in field isolates of Botrytis cinerea in Israel. Plant Pathol. 1992 41, 41–46.

26. Grabke A.: Fernández-Ortuño, D.: Schnabel, G.: Fenhexamid resistance in Botrytis cinerea from strawberry fields in the Carolinas is associated with four target gene mutations. Plant Dis. 2012 97, 271–276.

27. Jiang, J.: Ding, L.: Michailides, T. J.: Li, H.: Ma, Z: Molecular characterization of field azoxystrobin-resistant isolates of Botrytis cinerea. Pestic. Biochem. Physiol. 2009 93:72–76.

28. Fournier, E.: Giraud, T.: Loiseau, A.: Vautrin, D.: Estoup, A.: Solignac, M.: Cornuet, J.M.: Brygoo, Y.: Characterization of nine polymorphic microsatellite loci in the fungus Botrytis cinerea (Ascomycota).” Molecular Ecology Notes 2002 2(3): 253–255.

29. Fournier, E.: Gladieux, P.: Giraud, T.: Dr Jekyll and Mr Hyde fungus’: noble rot versus gray mold symptoms of Botrytis cinerea on grapes.” Evol Appl 2013 6(6): 960–969.

30. Diguta, C.F.: Rousseaux, S.: Weidmann, S.: Bretin, N.: Vincent, B.: Guilloux-Benatier, M.: Alexandre, H.: Development of a qPCR assay for specific quantification of Botrytis cinerea on grapes.” FEMS Microbiol Lett 2010 313(1): 81–87.

31. Suarez, M. B.: Walsh, K.: Boonham, N.: O’Neill, T.: Pearson, S.: Barker, I.: Development of realtime PCR (TaqMan) assays for the detection and quantification of Botrytis cinerea in planta. Plant Physiol Biochem, 2005 43, 890–9.

32. Schneider, C.A.: Rasband, W.S.: Eliceiri, K.W.: NIH Image to ImageJ: 25 years of Image Analysis Nat Methods 2012, 9(7), 671–675.

33. Huber, F.: Röckel, F.: Schwander, F.: Maul, E.: Eibach, R.: Cousins, P.: Töpfer, R.: A view into American grapevine history: Vitis vinifera cv. ’Sémillon’ is an ancestor of ’Catawba’ and ’Concord’. J. Grapevine Research 2016, 55, 53–56.

34. Backmann, L.: Wegmann-Herr, P.: Jürgens, A.: Scharfenberger-Schmeer, M.: Correspondence Affiliation, Neustadt, Rhineland-Palatinate, Germany. 2023, status (manuscript in preparation).

